# Automatic Tracing of Ultra-Volume of Neuronal Images

**DOI:** 10.1101/087726

**Authors:** Hanchuan Peng, Zhi Zhou, Erik Meijering, Ting Zhao, Giorgio A. Ascoli, Michael Hawrylycz

## Abstract

Despite substantial advancement in the automatic tracing of neurons' morphology in recent years, it is challenging to apply the existing algorithms to very large image datasets containing billions or more voxels. We introduce *UltraTracer*, a solution designed to extend any base neuron-tracing algorithm to be able to trace virtually unlimited data volumes. We applied this approach to neuron-tracing algorithms with completely different design principles and tested on challenging human and mouse neuron datasets that have hundreds of billions of voxels. Results indicate that *UltraTracer* is scalable, accurate, and about 3 to 6 times more efficient compared to other state-of-the-art approaches.

## Introduction

The three-dimensional (3-D) morphology of a neuron is crucial for establishing its connections and function in the context of brain circuits (Ascoli, 2015). Reconstruction of such neuron morphology from optical images is an important challenge in neuroscience (Acciai, et al, 2016). Substantial international efforts, e.g. the DIADEM competition (Liu, 2011) and the collaborative BigNeuron initiative (Peng, et al, 2015), have led to sizeable advances in this field. Yet, it remains an open and critical question how to effectively reconstruct, or trace, extremely large 3-D image volumes of long projection neurons having potentially complex arborization patterns.

Typically, due to the limited field of view of optical microscopy, the 3-D image volume of a large mammalian neuron, e.g. a pyramidal neuron, is produced using tiled scanning over the brain area where the neuron resides. When the voxels have sub-micron size in 3-D, the overall volume of such a neuron often amounts to tens of billions or even trillions of voxels. Most published neuron tracing methods to date were not designed to handle such a massive amount of data.

Here we introduce an intuitive, explorative method called *UltraTracer* to effectively trace virtually infinite 3-D image volumes. We extend *UltraTracer* to be a container of a variety of different base tracing algorithms that bear different design principles, enabling *UltraTracer* to aggregate the merits of previous methods. We have found *UltraTracer* suitable for reconstructing very large neuron morphology from a number of tests demonstrated below.

## Results

The core algorithm of *UltraTracer* (Figure 1) reconstructs a neuron structure as completely as possible from the available image data based on a formulation of maximum likelihood estimation (MLE). The underlying assumption is that the occurrence of a specific neuron structure could be modeled using the joint probability of all of its subparts given the image. Briefly, *UltraTracer* iteratively factorizes the joint probability based on progressive maximization of conditional probabilities of the occurrence of salient and continuous subparts of a neuron (Methods). Concretely, *UltraTracer* begins the tracing from a subarea around the soma of a neuron, typically a cube with at least 512^3^ voxels. The soma could be automatically detected using a previous method or manually determined by one computer-mouse click so it was not a limiting step (Methods). A base tracing method *T* is chosen from a pool of candidate tracing methods Ω. *UltraTracer* analyzes the reconstruction produced by *T* and detects the tips of the neuron (Figure 1B). All such tips are added to a tip-queue. Then all tips in the tip-queue are sorted by saliency in terms of their thickness, image-intensity, and continuity (Methods). Next, depending on the base tracing method *T*, *UltraTracer* automatically and adaptively defines a new subarea to trace (Figure 1B, referred to as the “tip-distribution based adaptive window” method, or TDAW), based on either the most prominent single tip, or a group of nearby tips on a polygonal face of the polyhedron of the already-traced image volume. The new reconstruction is merged onto the existing reconstruction. The already searched tips are then eliminated from the tip-queue, while new tips of the merged neuron reconstruction are added into the tip-queue. Subsequently, the tip-queue is sorted again based on saliency. The tracing procedure repeats until no new tips could be detected and the tip-queue is empty (Figure 1B). This way, *UltraTracer* is capable of exploring a virtually infinite volume of image by following where the neurite signal goes. The final neuron morphology is produced together with the radius estimation along the reconstruction (Figure 1C). In our implementation, we designed the software to quickly extract an arbitrary subvolume of interest from very large neuron image files (Methods). Therefore, *UltraTracer* can smoothly trace a massive image archive without the need to load a large amount of image voxels into computer memory.

**Figure 1.**
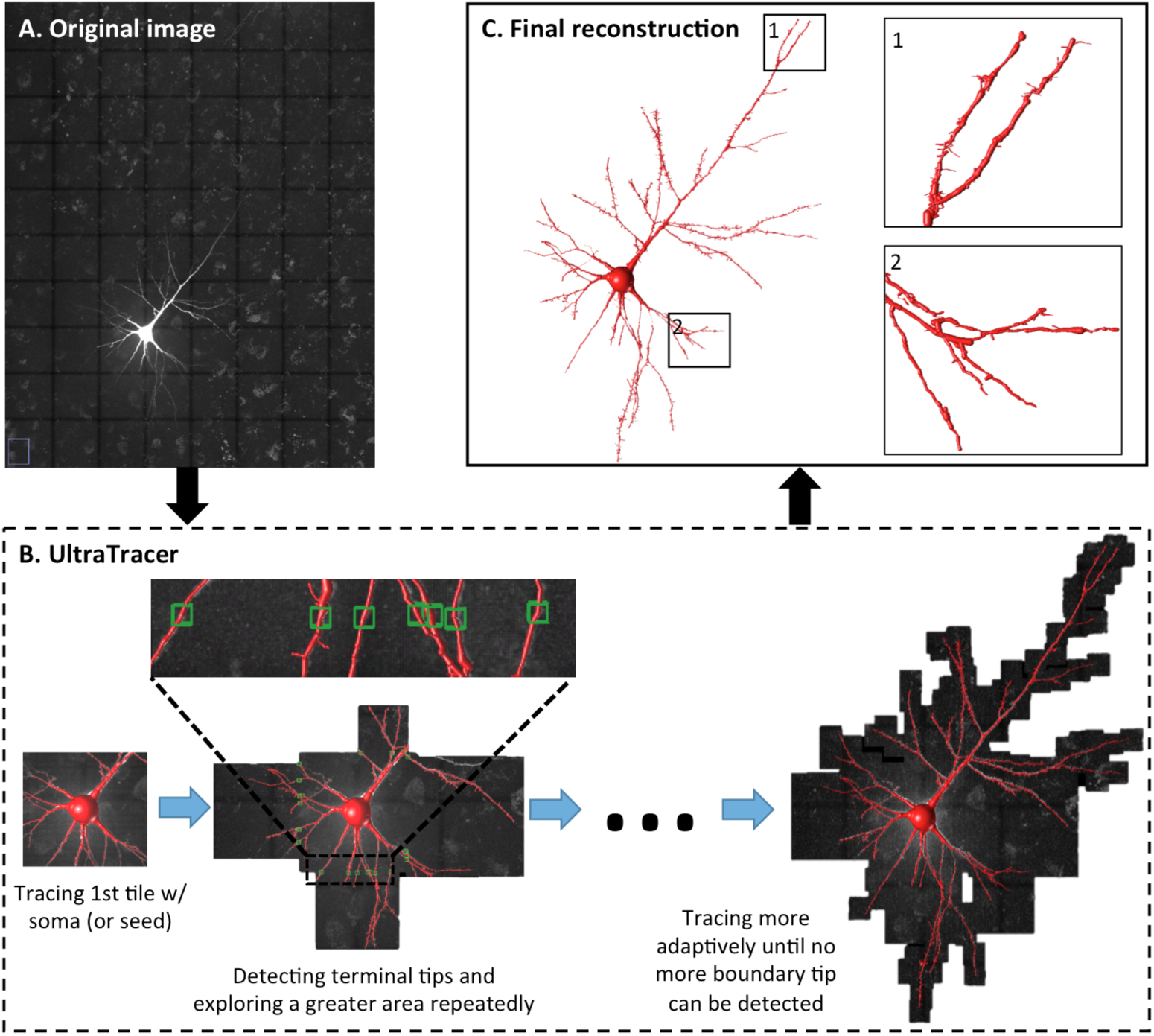
Workflow of UltraTracer for tracing a very large 3-D image volume. A. 3-D confocal image stack of a Lucifer Yellow labeled human pyramidal neuron. The voxel size is 0.18 × 0.18 × 0.5 µm and the overlaid grid (black lines) indicates how the image volume is subdivided into uniform tiles. B. UltraTracer first traces the subarea containing the soma and then detects the neuron terminal tips in the reconstruction and adaptively explores and traces neighboring subareas. Green boxes: terminal tips detected in tracing a subarea. C. Final reconstruction produced by UltraTracer with zooms of two parts for detailed visualization.

As a comprehensive test, in Ω we included four representative base tracing algorithms (Figure 2; Methods), namely APP1 (Peng, et al, 2011), APP2 (Xiao, et al, 2013), Neutube (Zhao, et al, 2011), and MOST (Wu, et al, 2014), available on the BigNeuron platform (Peng, et al, 2015; https://github.com/BigNeuron/BigNeuron-Wiki/wiki/Neuron-Reconstruction-Algorithms). These four methods have different design principles. They were also relatively robust, fast, and accurate compared to many other base tracing algorithms (Peng, et al, 2011; Xiao, et al, 2013; Zhao, et al, 2011; Wu, et al, 2014). Despite the differences between the outputs of these methods in tracing one single or multiple tree-shape neuronal arborization patterns from one single image tile, they can all be contained in the *UltraTracer* framework (Methods; Figure 2). Thus, *UltraTracer* extends arbitrary base tracing-algorithms to effectively trace across a very large image region adaptively (Figure 2A), a crucial utility that was not previously available to reconstruct massive scale datasets.

**Figure 2.**
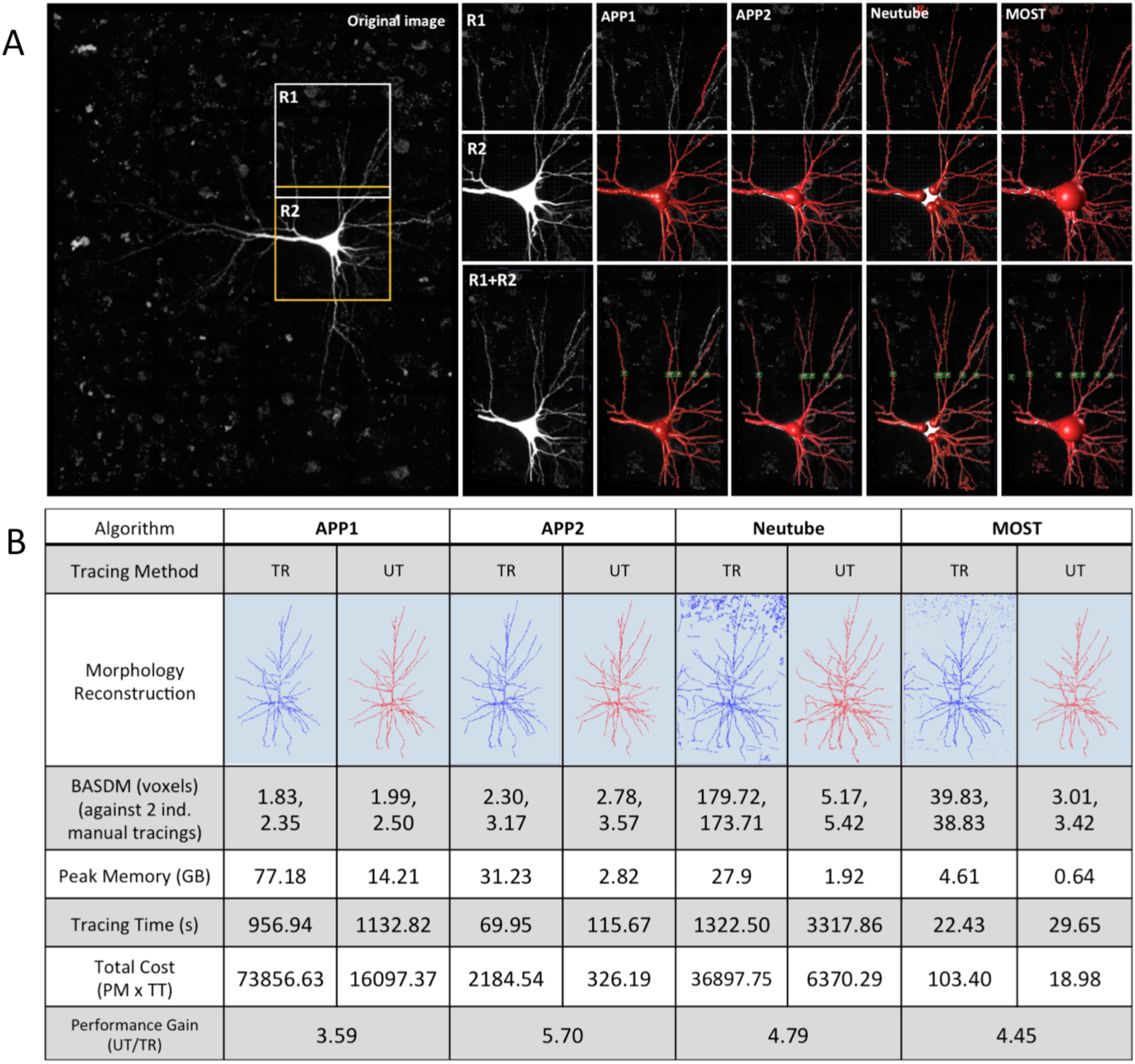
UltraTracer extends and improves various base tracers to reconstruct large image volumes. A. UltraTracer with four base tracers APP1, APP2, Neutube, and MOST (Methods) applied to image regions R1 and R2. B. Comparison of UltraTracer and the direct use of base tracers. TR: traditional method (i.e. using a base tracer directly to reconstruct the entire 3-D image volume); UT: UltraTracer; BASDM: Best Average Spatial Distance compared with Manual reconstructions; PM: Peak computer-Memory; TT: Tracing Time. The image volume used has 2111×3403×291 voxels. Two independent human manual reconstructions were used for comparison; their BASD (Best Average Spatial Distance) is 3.56 voxels.

Even for an image volume with about two billion voxels, which could still be handled by some base tracers directly, *UltraTracer* reduced dramatically the total amount of required computer-memory. The performance gain of *UltraTracer* over the direct use of certain base tracers was within the range of 3 to 6 times (Figure 2B). *UltraTracer* results were accurate as their average spatial distances to independent manual reconstructions were around 3 voxels, comparable to the spatial distance of the manual reconstructions themselves (3.56 voxels) (Figure 2B). In addition, for two base tracers, Neutube and MOST, *UltraTracer* had a gain of 10 to 30 folds in tracing accuracy (Figure 2B).

Since the best base tracer in Ω was APP2 in terms of speed and accuracy trade-off (Figure 2), we further tested the APP2-based *UltraTracer* on a series of images, of which the volume ranged from 0.47 to 521.5 billion voxels (Figure 3). *UltraTracer* was able to effectively trace only the sparse neuronal structures in these images, without spending time to analyze the entire data volumes (Figure 3). The data volume reduction in tracing was between about 3 and almost 40 times. Particularly, *UltraTracer* was the only automatic neuron tracing method that was applicable to ultra-volumes such as neurons 5, 6 and 7 that had 118, 122 and 521 billion voxels, respectively. The traditional approach also failed for neuron 4 because the actual peak-memory requirement to trace this dataset (14-billion voxels) exceeded the total amount of available memory (128GB) in our testing machine. When the image volume increased, we observed a bigger data-volume reduction rate in tracing. This matches well with expectation, since the neuron arborizations to be traced are roughly 1-D structures while the image data is 3-D, and thus the fraction of relevant space to be explored generally decreases with increasing data volume. The results indicated excellent robustness and scalability of *UltraTracer* for extremely large neurons. Of note, the accuracy of reconstructions produced by *UltraTracer* was similar to that of the conventional approach, when such a traditional approach was still feasible in our testing (Figure 3, Neurons 1, 2, and 3). Measured in terms of spatial distance, bifurcation points, and five other morphological and topological features, and compared against the statistics drawn from collections of reconstructions produced using control-images (Methods), the reconstructions produced by *UltraTracer* were consistent with those generated using the traditional approach when applicable (Figure 3, bottom-left).

**Figure 3.**
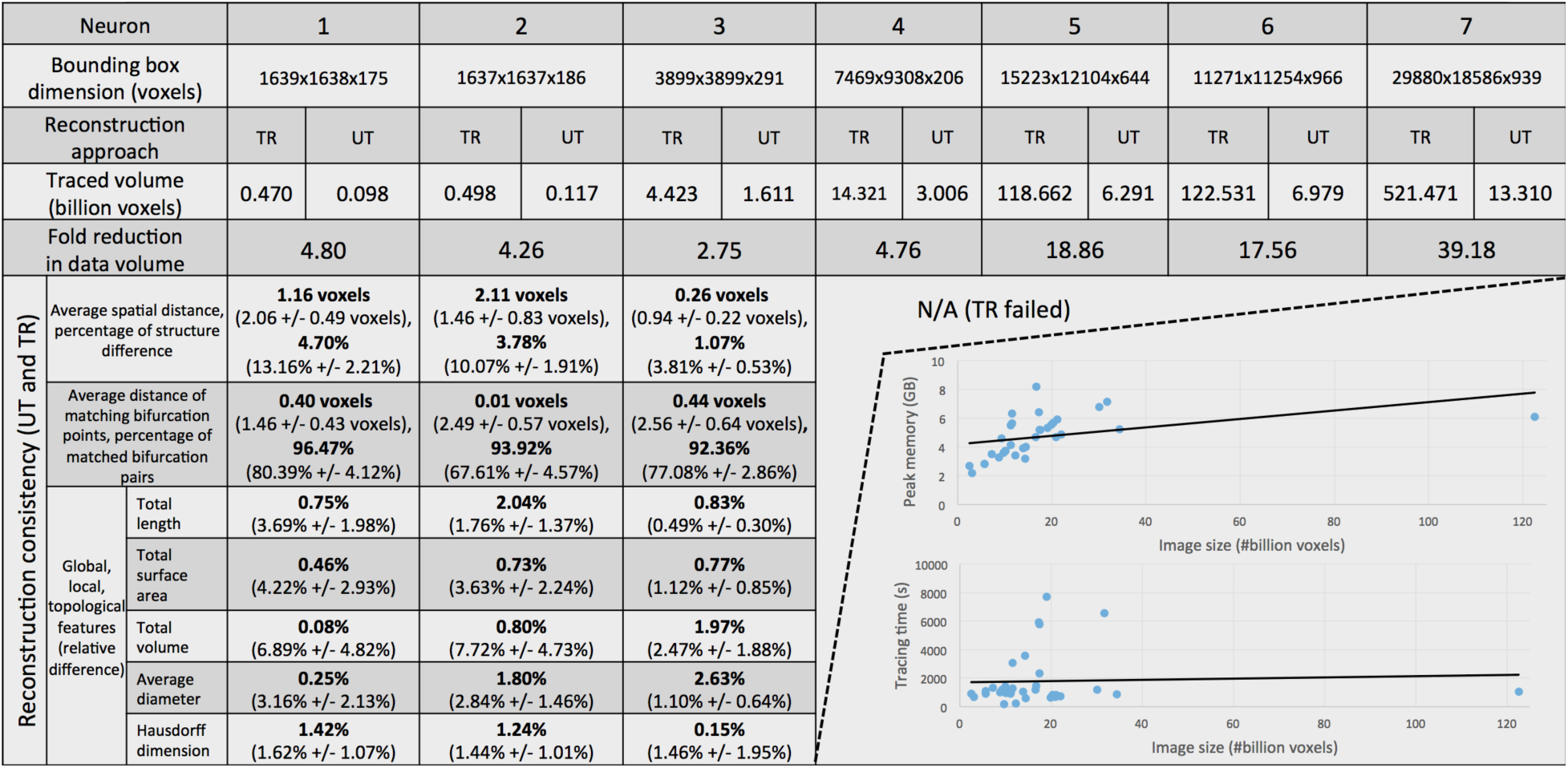
UltraTracer (with base tracer APP2) is scalable with respect to ultra-volumes of neuron images, without compromising the tracing accuracy in terms of spatial distance, morphological and topological features. TR: Traditional approach. UT: UltraTracer. Testing data: neurons 1, 2, 3, and 4 are confocal image stacks of human pyramidal neurons, neurons 5 and 7 are confocal image stacks of mouse pyramidal neurons, neuron 6 is a brightfield image stack of human pyramidal neuron. In reconstruction-consistency testing of TR and UT based on various features, the “percentage of structure difference” of two reconstructions measures the portion of their visible difference (the nearest matching reconstruction nodes in two tracings are more than 2-voxel apart), the “percentage of matched bifurcation pairs” is defined as the portion of reciprocally best matching bifurcation points divided by the average number of bifurcation points of two reconstructions, the “total length”, “total surface”, and “total volume” are the length, the surface and the volume of all neuronal compartments in reconstructions, the “average diameter” is the average diameters of all compartments in a reconstruction, the “Hausdorff dimension” (Falconer, 2004) measures the fractal dimension of reconstructions. In parentheses, the statistics (mean +/- s.d.) derived from TR-reconstructions using 59 rotated images (every 6 degrees around the center of XY-plane) for each neuron are shown as controls. Bottom-right inset: Regression analysis of peak memory and tracing time versus the image volume tested on 31 brightfield images.

In addition to confocal laser scanning images (Figures 2 and 3), we also tested *UltraTracer* using 31 challenging brightfield images of mouse and human neurons that had distinct appearance from laser scanning images (Figure 3, bottom-right insets; Supplementary Figure 1). After a number of tests we found these brightfield images were hard to trace successfully using the majority of automatic methods ported in BigNeuron. Differently, *UltraTracer* produced reconstructions that were consistent with visual inspection (Supplementary Figure 1). Both the peak memory and tracing time of *UltraTracer* scaled relatively smoothly on average with respect to the input image volume (Figure 3, insets).

Since *UltraTracer* was essentially a wrapper of any base neuron tracers, we also used it to combine multiple different base tracers (Supplementary Figure 2 and Supplementary Figure 3). For instance, we noted APP2 often traced well in the soma area while Neutube and MOST were sometimes more suitable to trace curvilinear structures. Thus in one variation of *UltraTracer*, we started the APP2-tracing for the image region around soma, followed by using Neutube or MOST for other image regions (Supplementary Figure 2). We found that in such a combination scheme, “APP2+Neutube” was able to explore 47% larger image area than “APP2+MOST” and thus the former generated a more visually complete reconstruction, even though the latter was 24 times faster than the former (Supplementary Figure 2). In a more complicated case, for every adaptively searched image region, we profiled the reconstructions generated by several base tracers. Then we chose either the best reconstruction or their consensus as the result from the current image region (Supplementary Figure 3). These variations of *UltraTracer* could be slower or faster than some of the base tracers (e.g. the “consensus” combination was slower than APP2 but faster than Neutube), and it could provide more consistent reconstructions compared to manual work (e.g. the combination reconstructions had more satisfaction in visual inspection as well as roughly 20% ~ 50% smaller distance scores than that of the Neutube results).

In addition to TDAW (Methods), we also considered using certain domain knowledge, or prior information, of neuron morphology to help refine the choice of the next tracing subarea. Our intuition was that a large window-size should be used for densely arborized image regions, and a small window-size would be sufficient for sparsely distributed neurites. Therefore, we estimated a lookup table of the average “expected” window size with respect to the distance between a neuron-compartment and its corresponding soma (Figure 4A), based on analyzing the spatial distribution of 968,348 neuron-compartments in 259 manually curated human and mouse neurons in the Allen Cell Types database (http://celltypes.brain-map.org/) and the BigNeuron initiative (Peng, et al, 2015) (Supplementary Figure 7) (Methods). Next, we used this lookup table as the prior information to guide TDAW (Methods). This new method, called the “prior-based TDAW” (PTDAW), enabled *UltraTracer* to trace human and mouse pyramidal neurons slightly more completely than TDAW (Figure 4B). Quantitatively, for the human neuron in Figure 2B, TDAW and PTDAW reconstructions were still close to each other (average spatial distance = 1.75 voxels). For a mouse pyramidal neuron (Supplementary Figure 4), we also observed similar performance of the two methods (average spatial distance = 2.85 voxels, comparable to the distances between each of these two reconstructions and the corresponding manual reconstruction, respectively 3.24 and 3.16 voxels).

**Figure 4.**
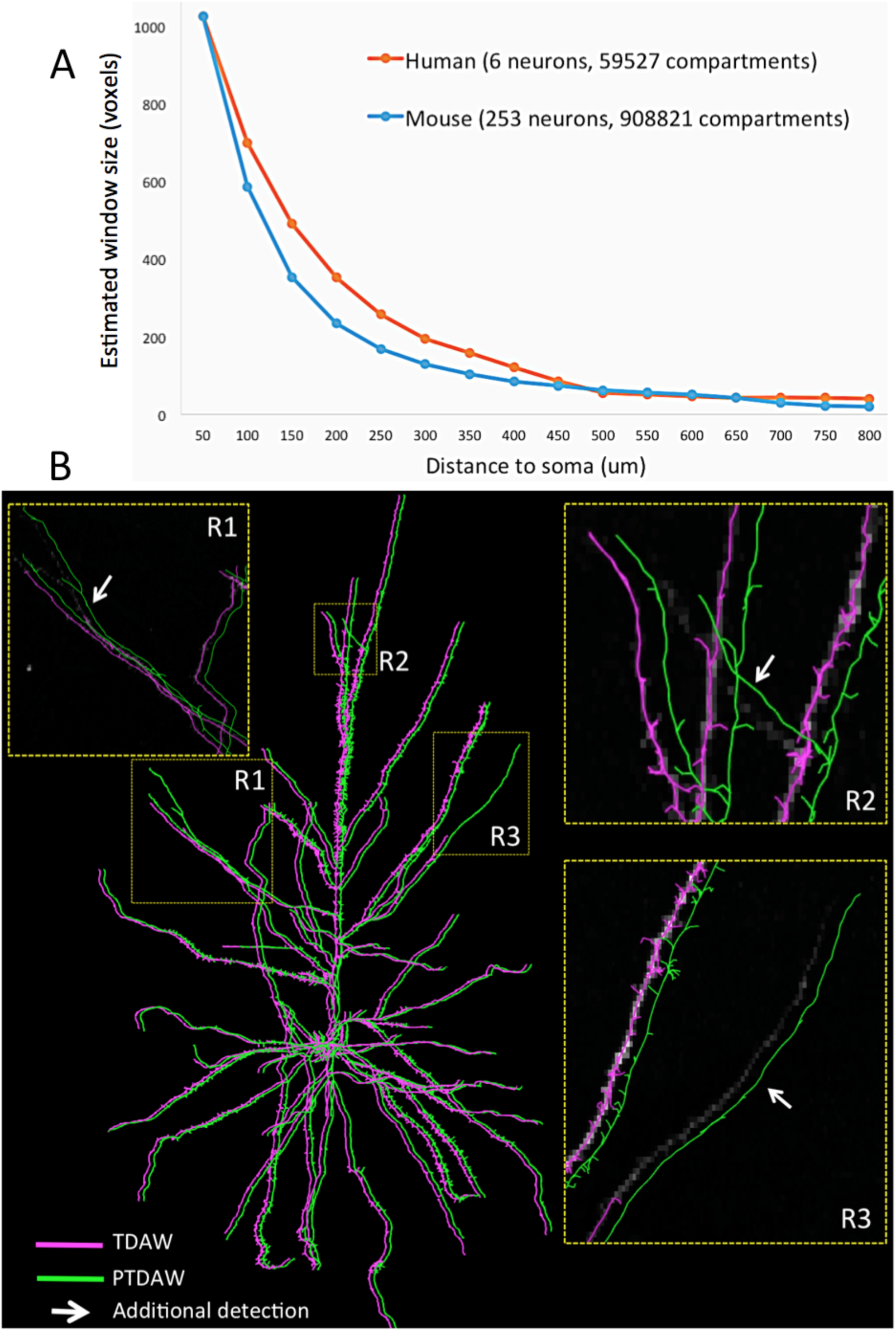
UltraTracer enhanced by incorporating prior knowledge of the adaptive subarea (window) size in tracing, which was learned from large-scale statistics of mammalian neuron reconstructions. A. The estimated window size (in x, y, and z) as a function of the distance of a neuron compartment to the soma. The maximum window size was set to be 1024 voxels. B. Comparison results of two tracings, one with the TDAW method (magenta) and another with PTDAW (green), where the prior is the estimated window size in A. In each zoom-in region (R1 ~ R3), the gray-scale image voxels are also displayed. The two reconstructions are offset slightly for better visualization.

Finally, instead of starting from a single soma location to trace one neuron, we also used *UltraTracer* to reconstruct human neurons, including their axons and dendrites, from separate but serial slices of brain tissue. We iteratively applied *UltraTracer* to multiple independent starting locations of the fragmented neuron structures, followed by stitching these fragments using Vaa3D (Peng et al, 2010) (Supplementary Figure 5). We manually validated one such example, which had totally 318.3 billion voxels in three separate sections. The total length of the tracing was 22.51mm. We found that about 90% of the compartments in the automatic reconstruction could be validated manually, while the other 10% were very challenging to reconstruct even for manual work.

## Acknowledgement

HP conceived this project, designed and managed this study, proposed the theoretical framework of the method, and wrote the paper with assistance from ZZ and other coauthors. ZZ developed the tip-queue based neuron growth algorithm, implemented the software and generated results, with the help of coauthors. We thank Allen Institute for Brain Science and data contributors of the BigNeuron project for providing neuron datasets. This work was funded by the Allen Institute for Brain Science. The authors wish to thank the Allen Institute founders, P. G. Allen and J. Allen, for their vision, encouragement and support.

## ONLINE METHODS

### *UltraTracer key* method and implementation

A neuron structure S can be modeled as the joint occurrence of its parts S_*i*_, *i*=1, …*N*. Given the image data D, the likelihood, i.e. the joint conditional probability, of occurrence of S is *L*(S|D) = *p*(S|D) = *p*(S_1_,S_2_, …, S_*N*_|D). The optimal neuron tracing problem can be formulated as a maximum likelihood estimation (MLE) problem, i.e. maximizing *L*(S|D) subject to constraints that the neuron’s parts should be maximally connected, and the connections should be as continuous, smooth, and biologically plausible as possible. This joint probability may be factorized in a combinatorial number of ways, depending on the definition, orders, and groupings of the neuron’s parts (substructures). Without loss of generality, max *L*(S|D) = max *p*(S_1_,S_2_, …, S_*N*_|D) = max *p*(S_2_, …, S_*N*_|D,S_1_)*p*(S_1_|D) = max *p*(S_*k*+1_, …, S_*N*_|D,S_1_,S_2_, …, S_*k*_)*p*(S_2_,…,S_*k*_|D,S_1_)*p*(S_1_|D). We used an intuitive approach to solve the MLE problem, by repeatedly finding the most probable substructures of S given the image. Obviously one such substructure should be the soma area of the neuron as well as immediately connected neurites. Then we iteratively detected other most probably connected substructures and grew the neuron reconstruction as completely as possible.

In our implementation, we first used the base tracer to reconstruct the cell body (soma) area. The size of the first subarea can be defined by the user, and the default size was 512 × 512 × 512 voxels. Then we designed a floating-search approach to smartly grow the neuron. Typically, the terminal tip close to the boundary indicates the continuity of the neuron structure. In order to assess where the neuron goes, all boundary tips from the previous tile’s reconstruction were detected as the reference locations. For single-root tracing (e.g. APP1 (Peng, et al. 2011) and APP2 (Xiao and Peng 2013)), each boundary tip was used as the root input to generate a single neuron tree on the adjacent tile. Different neuron trees starting from the previous tile’s boundary tip were continuously added to the adjacent tile. For multiple-segment tracing (e.g. Neutube (Zhao, et al. 2011) and MOST (Wu, et al. 2014)), all neuron structures on the adjacent tile were traced first by the algorithm. Within the reconstruction, only the segments containing the previous tile’s boundary tip were kept, and all other detected signals were removed. We also used 10% overlap with the adjacent tile to reduce the false negative rate. This floating-search approach not only solved the under-tracing problem from single-root tracing algorithms, but also eliminated the over-traced segments from multiple-segments tracing algorithms.

To efficiently explore the neuron structure, we used the density of boundary tips to adaptively define the next area. First, all possible boundary tips in all six directions (left, right, up, down, in, and out) were located. In each direction, all detected boundary tips were classified into different groups based on the neighbors’ distance. For each group in each direction, 1.2 × the maximum distance between two tips’ locations was defined as the x, y, and z dimensions of the next area. A minimum dimension (128 × 128 × 128 voxels) was predetermined in case the defined dimension was too small. With the adaptive window size, *UltraTracer* loaded a much smaller amount of image volume to reconstruct a neuron (Supplementary Figure 6). This method was called tip-distribution based adaptive window (TDAW). We also introduced a variant called “prior-based TDAW” (PTDAW). In PTDAW, we used Sholl analysis (Sholl, 1953) to collect the statistics of the neuron-compartment density with respect to the soma locations (Supplementary Figure 7). Then we converted the density per 3-D unit-volume to the expected window sizes in one dimension (assuming the size of a new tracing subarea to be the same in x, y, and z). Finally, in PTDAW, the new search window size was set to be the greater one of the window size estimated using TDAW alone and the respective window size value in the lookup table, but no larger than the size of the current window containing the border-tips of consideration. The latter constraint was specifically designed for pyramidal neurons, but could be relaxed for other types of neurons.

To avoid over-tracing or topological errors due to the overlap between adjacent tiles, we designed a simple fusion approach by calculating the overlap region between two reconstruction compartments from adjacent tiles. If it was greater than 50%, only the compartment from the first traced tile was kept. Otherwise, both compartments were considered to be valid (Supplementary Figure 8). All our reconstructions were represented by a number of compartments with ID, type, coordinates, radius, and parent information. When the base tracer did not provide useful radius information, we used a Vaa3D “neuron radius” plugin to calculate the radii.

The soma, as well as other potential seed locations for starting the tracing, was automatically detected using a gray-weighted distance-transform method (Xiao and Peng, 2013), or manually determined by the virtual-finger powered one computer-mouse click technique (Peng, et al, 2014).

### Computer configuration

We used a Linux machine with 8 Intel(R) Xeon(R) CPU E5-1620 0 @ 3.60GHz, 128 GB memory, and C++ programming language to calculate the computational cost including peak memory and tracing time.

### Software availability

*UltraTracer* is available in Vaa3D software (vaa3d.org), and it is open source (https://github.com/Vaa3D/vaa3d_tools/tree/master/hackathon/zhi/neurontracer). As long as the image format can be supported by Vaa3D, it can be explored in *UltraTracer*. However, for very large-scale images (> 100 billion voxels), the computer may not have enough memory to load the entire image. In that case, *UltraTracer* also supports several other image formats, specifically the Vaa3D-Terafly interface (Bria, et al, 2016) that includes 2D TIFF/Vaa3D raw files, single multipage 3D TIFF/Vaa3D raw file, three-leveled *y*-*x*-*z* hierarchy of tiles with 3D TIFF/Vaa3D raw files, and HDF5 volume.

## Supplementary Figures

**Supplementary Figure 1.**
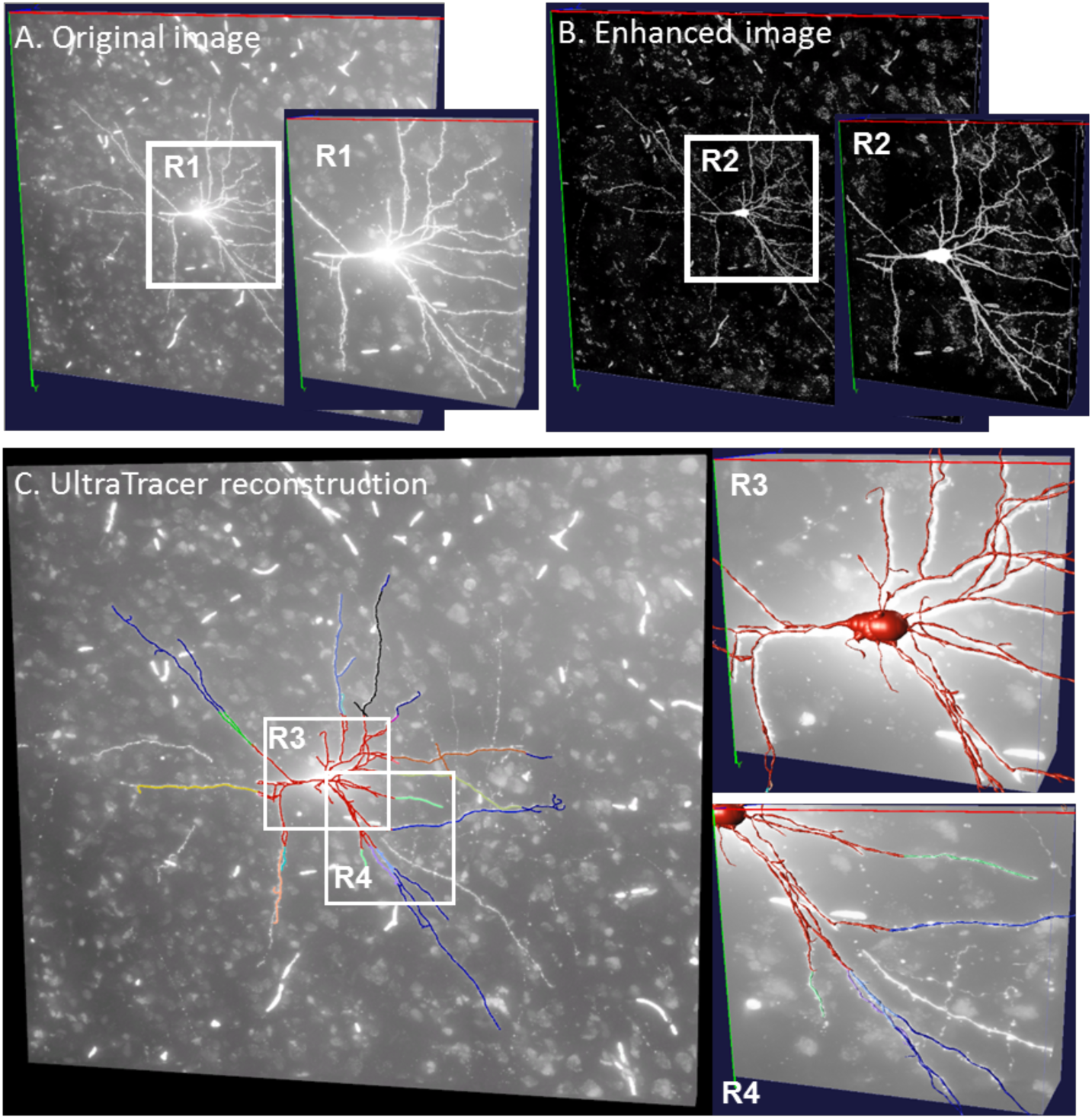
Application of UltraTracer to brightfield imaging image stacks of mouse V1 neurons. A. An example of brightfield image. B. Enhanced image using an adaptive approach (Zhou, et al, 2015). C. UltraTracer reconstruction based on the enhanced image in B. Different colors indicate reconstructions from different image regions.

**Supplementary Figure 2.**
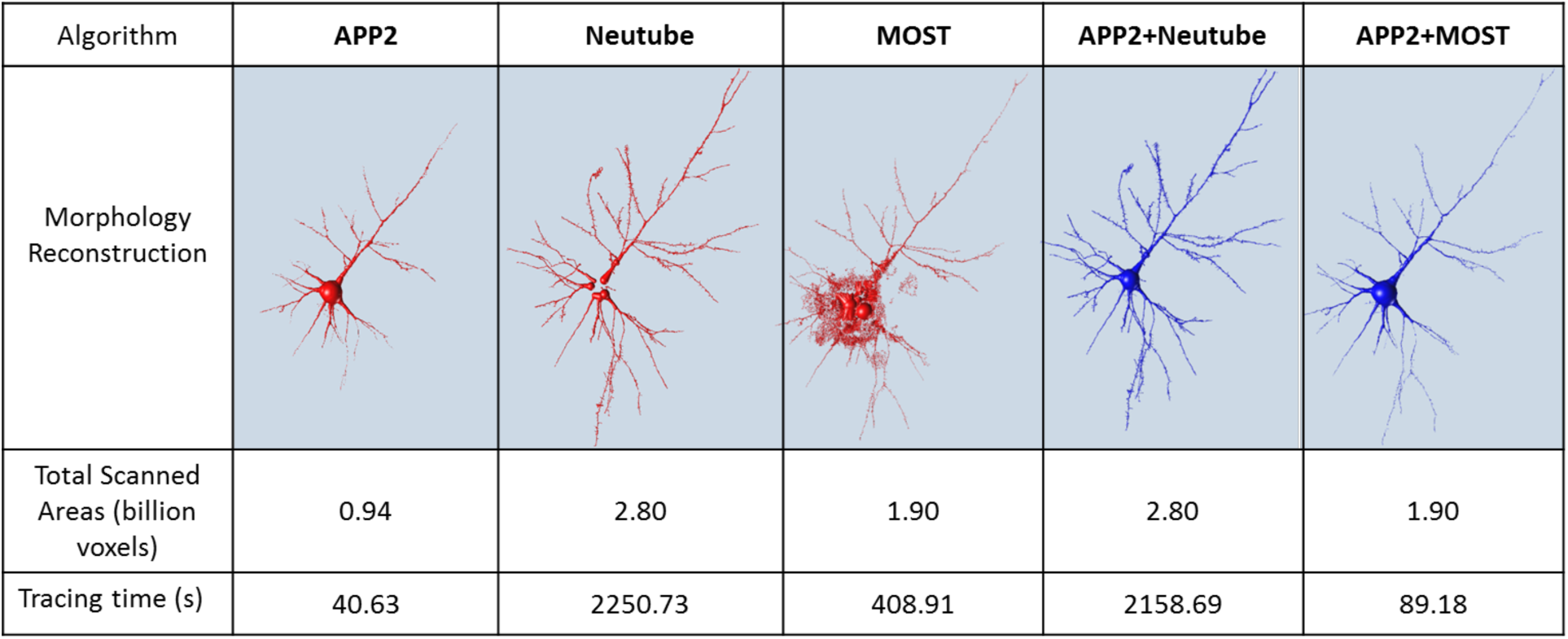
Combination scheme 1: UltraTracer combines different base tracers to achieve better performance on a 3-D confocal image stack of a Lucifer Yellow labeled human pyramidal neuron. APP2+Neutube: the soma region is traced by APP2, and the rest is traced by Neutube. APP2+MOST: the soma region is traced by APP2, and the rest is traced by MOST. APP2+Neutube explores 2.80 billion voxels areas, but needs 2158.69s for tracing. APP2+MOST generates a relatively complete reconstruction (1.90 billion voxels scanned areas) with a much faster tracing speed (89.18s tracing time). Neuron data used here is the neuron 4 in Figure 3.

**Supplementary Figure 3.**
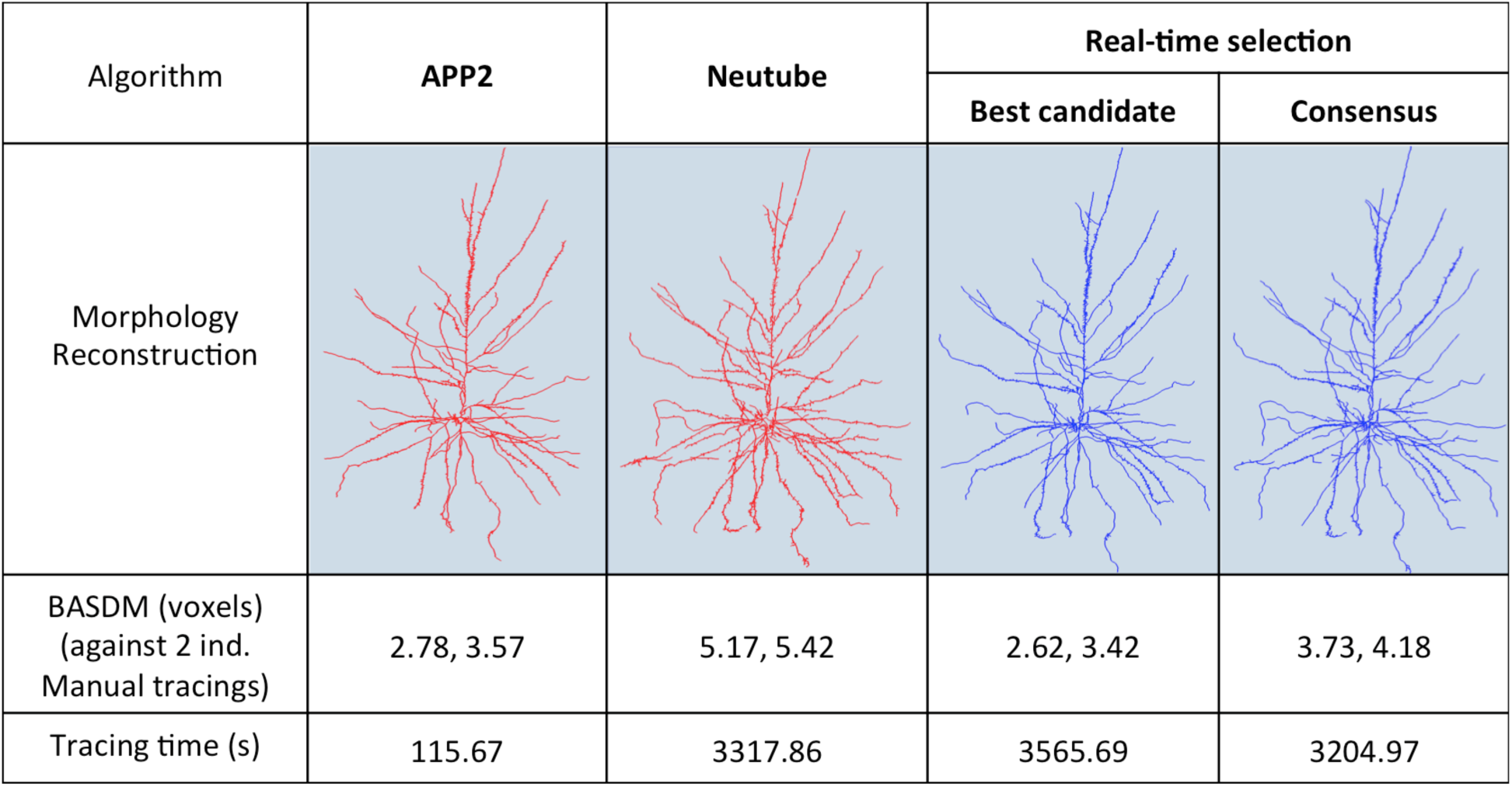
Combination scheme 2: UltraTracer real-time selects suitable tracing algorithm on a confocal image stack of human pyramidal neuron. For each explored image region, two reconstructions (APP2 and Neutube) were generated first. For the “best candidate” result, the contrast-to-background ratio in the image region around the reconstruction was used to choose the suitable algorithm. For the “consensus” result, the union of two reconstructions is used as the result for the current image region. Both two real-time selection results had similar BASDM scores (2.62 voxels and 3.42 voxels in the best candidate result, and 3.73 voxels and 4.18 voxels in the consensus result) to APP2 (2.78 voxels and 3.57 voxels) and Neutube (5.17 voxels and 5.42 voxels). Neuron data used is the same neuron in Figure 2B.

**Supplementary Figure 4.**
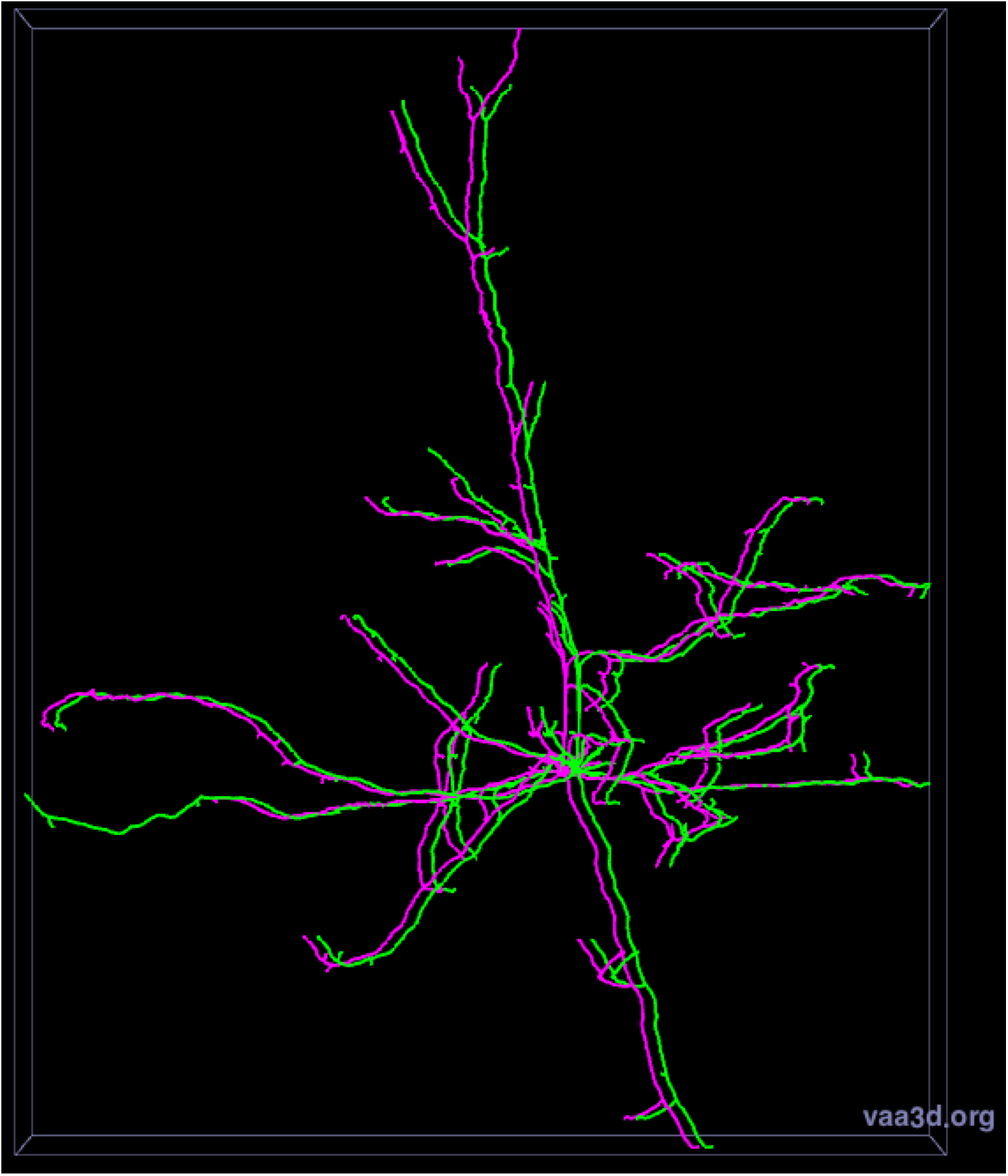
Tracing results of TDAW (magenta) and PTDAW (green) for a mouse pyramidal cell (voxel size 0.143µm×0.143µm×0.28µm).

**Supplementary Figure 5.**
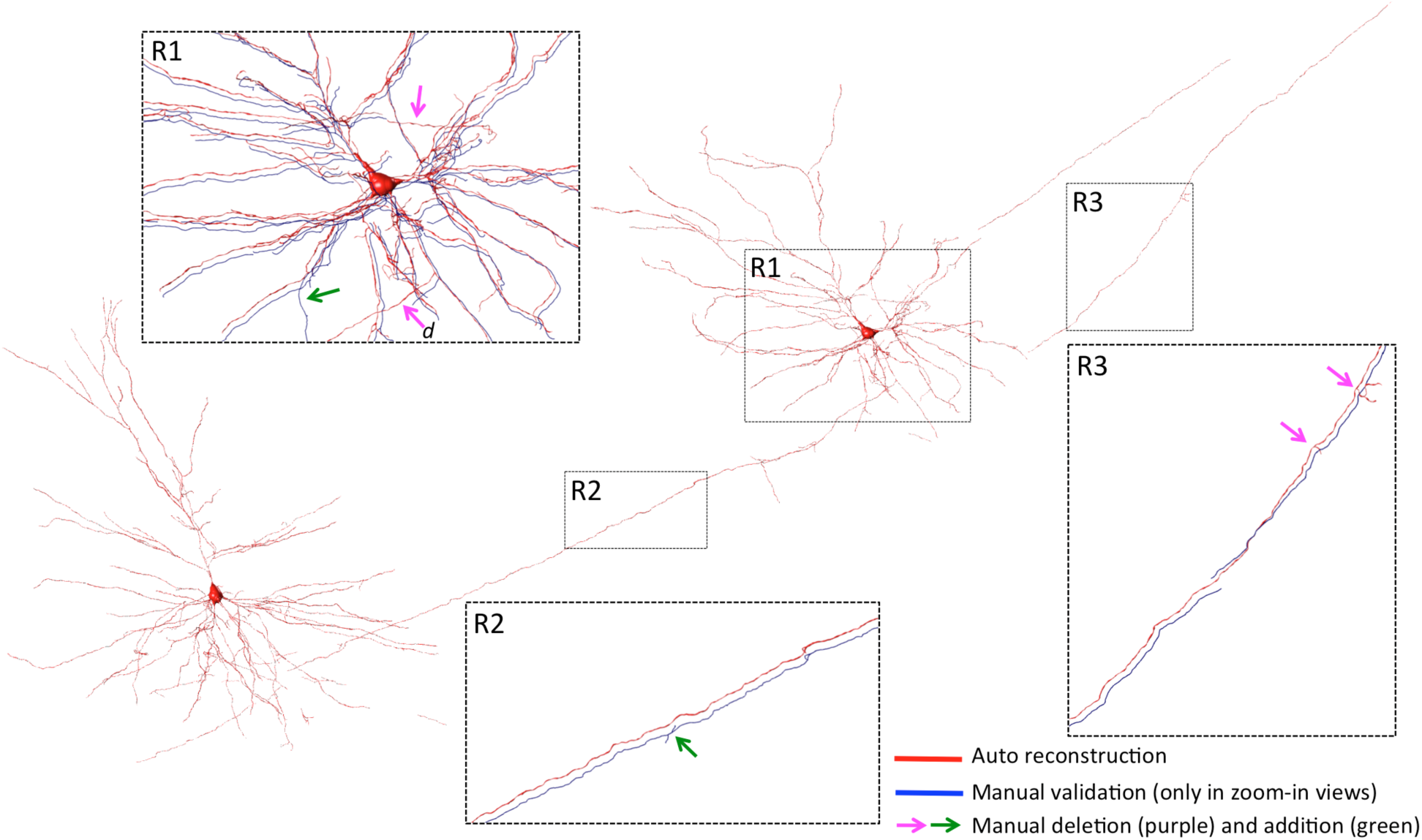
An application example of UltraTracer for tracing multiple biocytin-filled human neurons with axons. The images were from 3 sections, each of which was imaged separately (voxel size 0.114µm×0.114µm× 0.28µm). UltraTracer was used to reconstruct automatically based on multiple starting locations on these separate image stacks. The final reconstruction (red) was assembled using the NeuronAssembler tool in Vaa3D (vaa3d.org). The reconstruction, including axons and dendrites, was also manually validated (blue in zoom-in views, slightly offset for better visibility), with some substructures of the reconstruction edited (addition or deletion of some structures based on visual inspection). Overall more than 90% portion of the automatic reconstruction could be easily validated manually for this example, while the 10% were too difficult even for manual reconstruction (e.g. the manual deletion in location d of region R1 seemed to be a problematic deletion in the manual correction). The total lengths of the automatic and manually curated reconstructions were 22.51 and 20.15 mm, respectively.

**Supplementary Figure 6.**
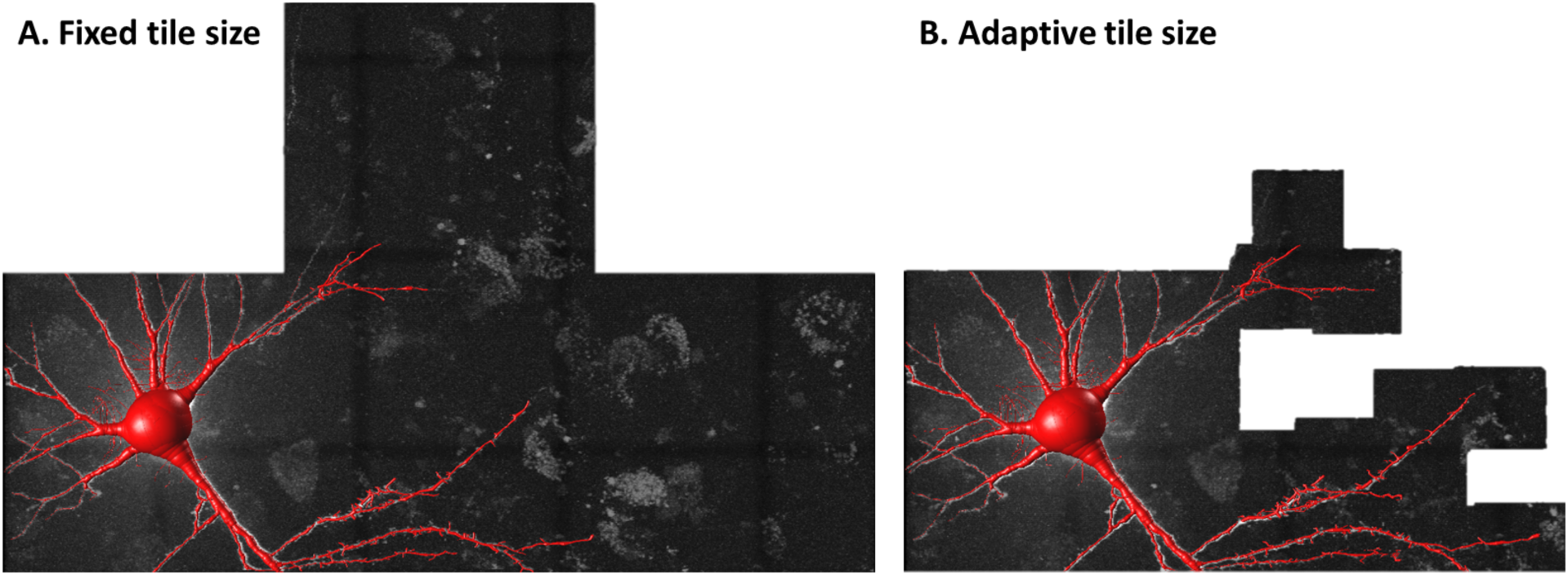
Fixed versus adaptive tile size. A. Based on the boundary tips of the left tile (containing the soma), three image tiles (1.1 billion voxels) have been loaded to trace the right side of the neuron with fixed tile size. B. A much smaller part (0.55 billion voxels) of the image volume has been loaded with the adaptive tile size.

**Supplementary Figure 7.**
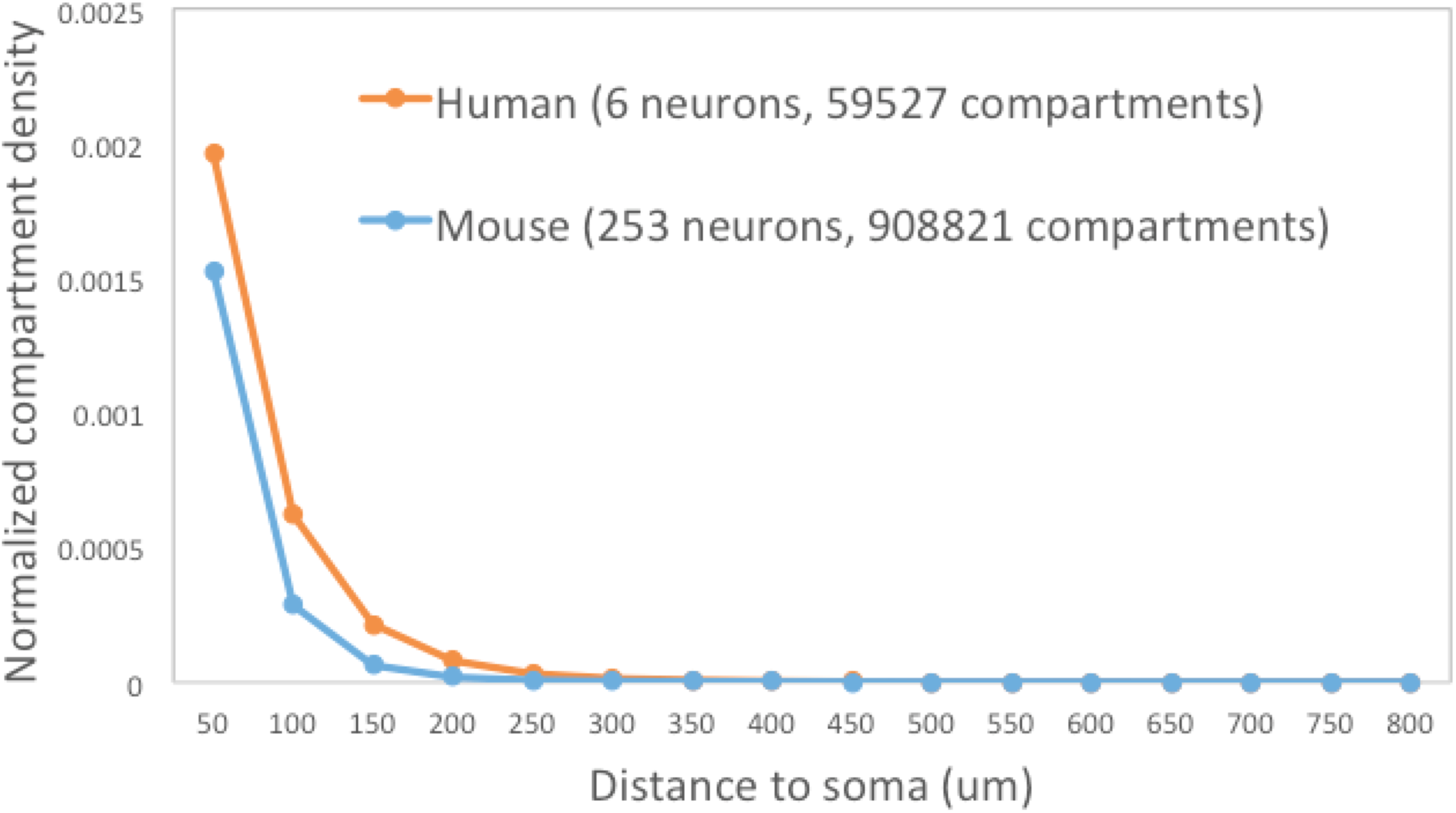
Average neuron-compartment density as a function of the distance between the neuron-compartment and the soma.

**Supplementary Figure 8.**
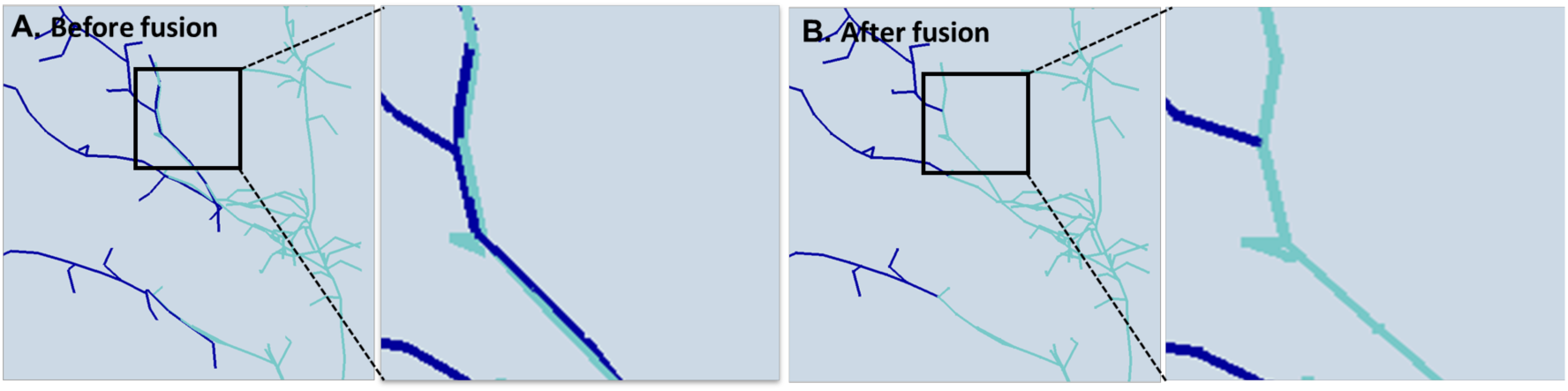
One reconstruction fusion example. A. Over-tracing due to overlap between adjacent tiles. B. The over-tracing error has been fixed.

